# Rapid detection of Pecan Root-Knot Nematode, *Meloidogyne partityla*, in laboratory and field conditions using loop-mediated isothermal amplification

**DOI:** 10.1101/2020.01.09.900076

**Authors:** Sumyya Waliullah, Jessica Bell, Tammy Stackhouse, Ganpati Jagdale, Abolfazl Hajihassani, Timothy Brenneman, Md Emran Ali

**Affiliations:** University of Georgia, Department of Plant Pathology, Tifton, GA 31793, USA; University of Georgia, Department of Plant Pathology, Athens, GA 30602, USA

**Keywords:** Pecan, LAMP assay, Root-Knot Nematode, *Meloidogyne partityla*, molecular diagnosis

## Abstract

*Meloidogyne partityla* is the dominant root-knot nematode (RKN) species parasitizing pecan in Georgia. This species is known to cause a reduction in root growth and a decline in yields from mature pecan trees. Rapid and accurate diagnosis of this RKN is required to control this nematode disease and reduce losses in pecan production. In this study, a loop-mediated isothermal amplification (LAMP) method was developed for simple, rapid and on-site detection of *M. partityla* in infested plant roots and validated to detect the nematode in laboratory and field conditions. Specific primers were designed based on the sequence distinction of internal transcribed spacer (ITS)-18S/5.8S ribosomal RNA gene between *M. partityla* and other *Meloidogyne* spp. The LAMP detection technique could detect the presence of *M. partityla* genomic DNA at a concentration as low as 1 pg, and no cross reactivity was found with DNA from other major RKN species such as *M. javanica*, *M. incognita* and *M. arenaria*, and *M. hapla*. We also conducted a traditional morphology-based diagnostic assay and conventional polymerase chain reaction (PCR) assay to determine which of these techniques was less time consuming, more sensitive, and convenient to use in the field. The LAMP assay provided more rapid results, amplifying the target nematode species in less than 60 min at 65°C, with results 100 times more sensitive than conventional PCR (~2-3 hrs). Morphology-based, traditional diagnosis was highly time-consuming (2 days) and more laborious than conventional PCR and LAMP assays. These features greatly simplified the operating procedure and made the assay a powerful tool for rapid, on-site detection of pecan RKN, *M. partityla*. The LAMP assay will facilitate accurate pecan nematode diagnosis in the field and contribute to the management of the pathogen.

## Introduction

Pecan (*Carya illinoinensis*) is an important nut crop in North America. A variety of diseases caused by bacteria, fungi, virus, and nematode can attack these trees, and if not properly managed, they can cause economic damage to pecans. Root-knot nematodes (RKNs) of the genus *Meloidogyne* are economically important plant-parasitic nematodes, which cause significant damage to pecan production. Three species of RKNs, *Meloidogyne incognita*, *M. arenaria* and *M. partityla* have, have been reported as pathogenic to pecan [1–3]. Among these species *M. partityla* is the dominant RKN parasitizing pecan which has a greater incidence in southern United States [2–6]. *M. partityla* was originally found infecting pecan in South Africa, and was likely introduced into the United States during importation of infected pecan seedlings [7]. Due to the complexity and lengthy process of diagnosis and species identification, it is difficult to conduct extensive surveys, which are required for determining the pecan RKN distribution. Therefore, a quick diagnosis approach will be helpful to know the latest status of the pecan RKNs in southern United States.

Detection of the RKNs based on the morphological characteristics is an extremely difficult diagnostic task due to the RKN small, microscopic size and the difficulty distinguishing key diagnostic characters/features under a conventional light microscope [8–10]. In addition, the highly conserved and identical morphology across of *Meloidogyne* spp. make it even harder for their accurate identification [11]. Besides morphological characteristics, RKNs can also be identified based on isozyme patterns, and host plant response to infection, which are is also challenging for the identification at the species level [8, 12]. Furthermore, extensive knowledge of nematode taxonomy is required for proper RKN identification. Nucleic acid-based molecular methods mostly rely on the polymerase chain reaction (PCR), such as species-specific PCR, multiplex PCR, real-time PCR (qPCR), and PCR-fragment length polymorphism (PCR-RFLP) [13, 14]. Furthermore, random amplified polymorphic DNA (RAPD), high throughput genome sequencing, DNA microarrays, and satellite DNA probes-based hybridization techniques are also been reported to distinguish nematode species [11, 15, 16]. However, all of these molecular methods are time-consuming and require access to sophisticated and bulky laboratory equipment. In particular, for *M. partityla* identification, a PCR based assay targeting the fragment between mitochondrial COII gene and the large (16S) rRNA gene of mitochondrial DNA regions is widely used. It is a fast and sensitive tool compared to existing traditional methods for identification [13]. However, this PCR based diagnostic procedure still needs more time (~2-3 hrs) and expensive laboratory instrumentation. Due to these drawbacks, a detection method that is not only quick but also simple and economical is clearly needed.

Loop-mediated isothermal amplification (LAMP) is a novel technique that can overcome many of the limitations of traditional microscopy and molecular PCR based diagnostic assays [17–20]. LAMP has shown that the sensitivity can be 100 to 1,000 times higher than conventional methods and can easily detect below 1pg/µl or lower concentration [21]. This method costs less time per sample and is simpler to perform than other detection methods. It can be carried out rapidly (often in 30 min) with minimal equipment (a water bath or isothermal heat block) which is highly applicable for the onsite diagnosis of pathogens [22, 23]. Recently LAMP technique has been used for identifying different species of RKNs including *M. enterolobii*, *M. incognita*, *M. arenaria*, *M. javanica*, *M. hapla*, *M. chitwoodi* and *M. fallax* [22, 24, 25]. However, so far there is no report for using LAMP technique for the identification of *M. partityla*.

Due to the limitations of currently used methods, our objective was to develop a loop-mediated isothermal amplification (LAMP) assay for specific and rapid detection of *M. partityla* in pecan under laboratory and field conditions, and to determine the time needed to conduct the procedure.

## Materials and methods

### Source of Root-knot Nematodes

*Meloidogyne* spp. and all other plant nematodes used in this study were collected from the Nematode Laboratories, University of Georgia. All *Meloidogyne* spp. were purified from a single egg-mass. *M. partityla* was collected from infested roots of a pecan tree whereas egg-masses of each of *M. hapla*, *M. javanica*, *M. incognita* and *M. arenaria* were collected from infested roots of tomato plants maintained in the greenhouse in Athens, GA.

### Morphological assay

For morphological identification, eggs of *M. partityla* were collected from RKN infested pecan roots following method of Hussey and Barker (1973) and the species identification was performed following the protocol as mentioned in S1 Fig. In brief, after collecting adult females of pecan RKN, perineal patterns were prepared for the species identification. The collected RKNs were also transferred to a petri dish containing water and allowed to hatch into second-stage juveniles (J2) for 7 days at room temperature (25 °C). For morphological measurements, 20 J2 were hand-picked and temporary water mounts were prepared. Morphological measurements of each J2 were recorded with a Leica DME compound reseach microscope using an eyepiece micrometer at 400x magnification.

### Laboratory DNA extraction

Two different methods were used to extract genomic DNA from the root knot nematode. For the laboratory optimization step, genomic DNA was extracted from an adult female of each RKN species using the QIAGEN DNeasy® Blood & Tissue Kit (Qiagen, Valencia, CA) with some modifications. Briefly, the adult female was punctured with a dissecting needle 6–8 times, and added in 90 µl ATL buffer and incubated overnight at 56°C after adding 20 µl of proteinase K. The female was then vortexed for 15 sec before adding 100 µl Buffer AL and 100 µl ethanol (96%). The mixture was placed in a DNeasy mini spin column and centrifuged at 8000 rpm for 1 min and flow-through was discarded. With a new receiving tube, the centrifugation procedure was repeated adding 200 µl Buffer AW1 and then 200 µl Buffer AW2 to dry the DNeasy membrane at 14000 rpm for 3 min, discarding the flow through each time. The DNeasy Mini spin column was placed onto a 1.5 ml microcentrifuge tube and 50 µl Buffer AE was directly pipetted onto the DNeasy membrane to elute the DNA. Then the DNA was allowed to stand for 15 min before being centrifuged at 6000 rpm for 4 min. The resulting DNA was stored at –20°C for subsequent experiments.

### PCR amplification

The forward primer C2F3 and the reverse primer 1108 were used to amplify the fragment between the mitochondrial COII gene and the large (16S) rRNA gene [13]. The amplification was carried out under the following cycling conditions: 94 °C for 5 min, then 35 PCR cycles of 94 °C for 30 seconds, 55 °C for 30 seconds, 72 °C for 30 seconds and final incubation at 72 °C for 10 min. PCR products were analyzed by electrophoresis on a 1% agarose gel and imaging them in a UV gel doc machine. This primer (C2F3/1108) was used to detect the sensitivity of traditional PCR. Forward and reverse primer pairs are listed in S1 Table.

### Design of LAMP primers

LAMP primers were designed based on the sequences of the internal transcribed spacer (ITS) of the ribosomal DNA using PrimerExplorer V5 software1 (Eiken Chemical Co., Tokyo, Japan). Six primers were constructed: two outer primers (F3 and B3), a forward inner primer (FIP), a backward inner primer (BIP), a loop forward primer (LF) and a loop backward primer (LB). FIP comprised the F1c sequence complementary to the F1 and F2 sequence. BIP consisted of the B1c sequence complementary to the B1 and B2 sequences (Fig 1 and S1 Table).

**FIGURE 1.**
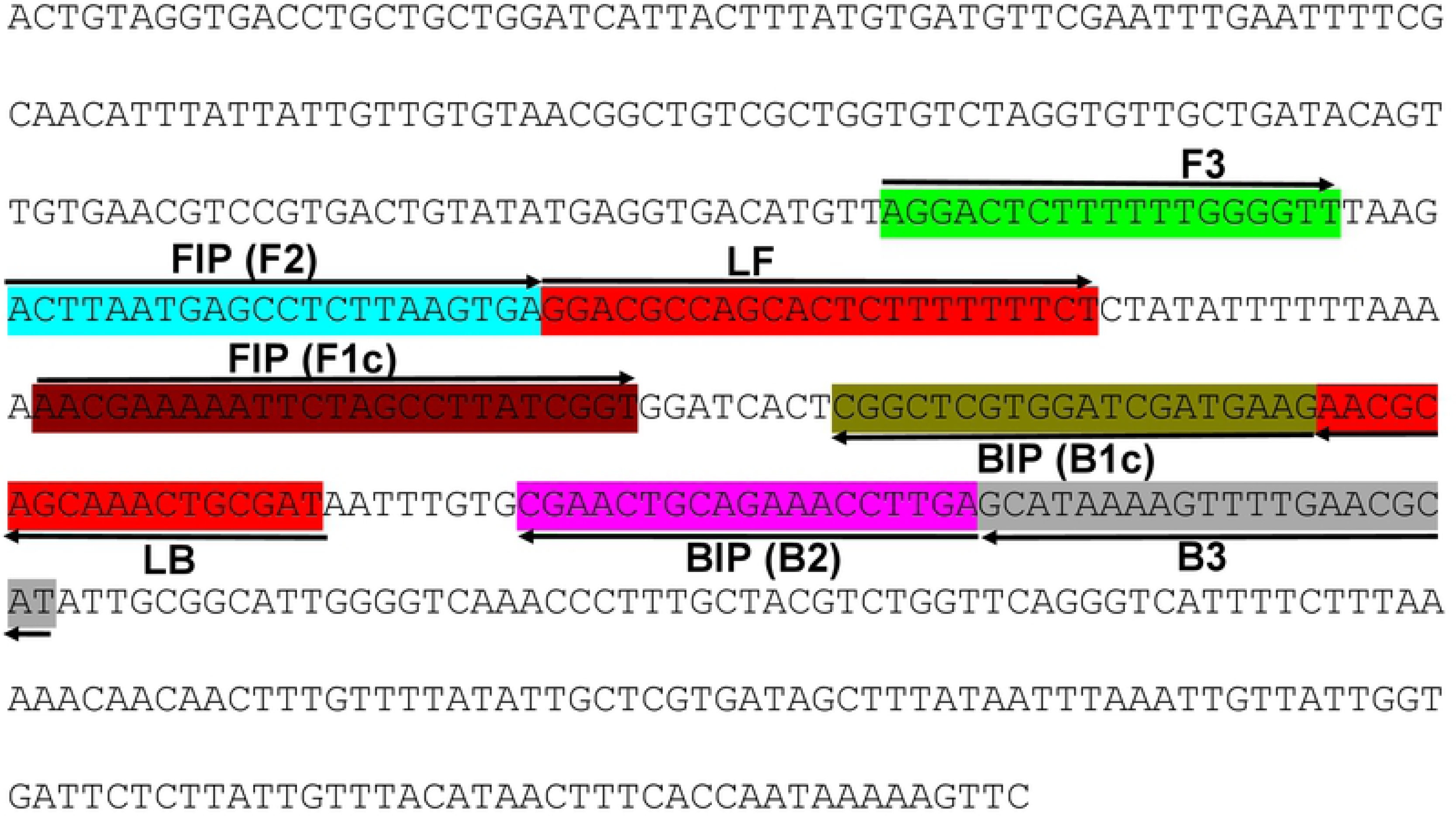
Location and partial sequence of Loop-mediated isothermal amplification (LAMP) primer sets targeting *Meloidogyne partityla* specific gene internal transcribed spacer (ITS)-18S/5.8S ribosomal RNA. Primer locations for LAMP assay (F3, B3, FIP [F1c-F2] and BIP [B1c-B2]). FIP is a hybrid primer consisting of the F1c sequence and the F2 sequence. BIP is a hybrid primer consisting of the B1c sequence and the B2 sequence. Arrows indicate the extension direction.

### Optimization of the LAMP reaction

The LAMP reaction was performed using LavaLAMP™ DNA Master Mix (Lucigen, WI, USA) according to manufacturer’s instructions. Each reaction contained 12.5 μl 2X master mix, 2.5 μl of primer mix, 1 μl DNA samples and the rest were filled with DNase/RNase free PCR certified water (TEKNOVA, Hollister, USA) to a final volume of 25 μl. The LAMP assay was optimized using different LAMP primer concentrations [F3, B3 (0.1–0.5 µM each); LF, LB (0.5–1.0 µM each) and FIP, BIP (0.8–2.4 µM each)], durations (30–60 min), and temperatures (65–73 °C). Optimization of temperature was performed in the Genie® III real-time instrument by determining the value of two parameters: amplification time (Tiamp) and amplicon annealing temperature (Ta). The temperature with least amplification time (Tiamp) and highest peak value of the melting curves (Ta) was considered as the optimum temperature for the assay. After optimization, all reactions were performed in 0.2-ml micro-tubes in a thermocycler (or Genie® III) set to 70°C for 60 minutes and terminated by incubating at 4°C for 5 minutes or for Genie® III amplification, the mixture was preheated at 90°C for 3 min, amplified at 70°C for 60 min, and then terminated at a range from 98 °C to 80 °C, with a decline rate of 0.05 °C per second. Each run contained a positive control of pure culture DNA and a negative control of PCR grade water without template.

### Analysis of LAMP products in laboratory

The LAMP amplification results were detected with three different methods in a laboratory settings: 1. on 1% TBE agarose gel stained with GelGreen® Nucleic Acid Stain (Biotium, Fremont, CA) for visual observation; 2. with SYBR™ Green 1 nucleic acid gel stain (Invitrogen, Carlsbad, CA) for observation under UV light; and 3. using Genie® III (OptiGene, Horsham, WS, UK) instrument by obtaining the amplified curves and analyzing data using Genie Explorer software (OptiGene, Horsham, WS, UK). All reactions were repeated at least three times.

### Sensitivity comparison of LAMP to conventional PCR

To determine the sensitivity of the LAMP assay, extracted DNA concentration was measured using a NanoDrop Lite (Thermo Scientific) instrument, and a tenfold serial dilution from the extracted DNA was done from 100 ng/μl down to a concentration of 10^−5^ ng/μl and subjected to the LAMP assay and the conventional PCR. Sensitivity tests were repeated three times.

### Specificity of the LAMP assay

Genomic DNA isolated from several *Meloidogyne* spp. including *M. partityla, M. hapla*, *M. javanica*, *M. incognita* and *M. arenaria* was used to verify the specificity of the LAMP assay. Specificity tests were repeated three times.

### On-site detection of *M. partityla*

For the onsite quick diagnosis application, suspected pecan root samples were collected from the UGA Ponder research farm pecan orchard in Tift County, GA, washed with distilled water, and examined under a handheld magnifier (10x) glass for presence of galls/knots (SPI Swiss Precision Instrument Inc., CA, USA). A single adult female was isolated from a gall and transferred into a 1.5 ml microcentrifuge tube, and DNA was extracted at the field using quick Extract-N-Amp™ Tissue PCR Kit (Sigma-Aldrich Co. LLC, MO, Louis. St; USA) with some modifications. Briefly, the single female of RKN was punctured with a dissecting needle 6–8 times and added into a tube containg 50 μl of extraction buffer and 12.5 µL of tissue preparation solution, which was mixed by pipetting up and down for several times. After initial incubation at room temperature for 10 minutes, sample was further incubated at 65 °C for 3 minutes. Finally, 100 µL of neutralization solution was added to the tube and mixed it by vortexing and 5 μl of extraction product was directly used as a template for LAMP amplification (S2 Fig). WarmStart® Colorimetric LAMP 2X Master Mix (New England Bio Labs Ltd., UK) which utilizes pH sensitive phenol red as an end-point indicator was used for onsite detection according to manufacturer’s instruction. Detection was carried out by observing color change of the WarmStart® colorimetric dye in the naked eye for the field detection. If a Genie® III instrument was used for the onsite detection, the amplification graph was observed on the screen to determine the result.

## Results

### Morphological identification

Based on the morphological characteristics the tested RKN’s were diagnosed as *M. partityla*. Morphological measurements (mean ± SD, range) of RKN J2 (n= 20) included mean body length (441.56 ± 17.41 µm, 412.50–468.75 µm), maximum body width (15.89 ± 1.22 µm, 14.06–18.75 µm), stylet length (14.76 ± 1.26 µm, 11.71-15.62 µm), tail length (48.20 ± 2.45 µm, 43.75–53.13 µm)), and hyaline region of the tail (15.82 ± 0.94 µm, 14.06–18.75 µm).

### Optimization of LAMP reaction

The LAMP assay was optimized using different LAMP primer concentrations, incubation temperatures and duration using DNA extracted from *M. partityla*. The final LAMP primer mix used in this study was: 0.2µM of each F3 and B3 primers, 0.8µM of each Loop F and Loop B primers and 1.6µM of each FIP and BIP primers (data not shown). For incubation temperature and time, the best reaction performance was found 70 °C for 60 minutes in 0.2-ml micro-tubes in a thermocycler or Genie® III (Fig 2, S2 Table). After gradient amplifications for the optimization of temperature, the LAMP products presented characteristic ladder-shaped band pattern when separated using 1% agarose gel electrophoresis (Fig 2A), where brightest band was obtained from 70 °C (S2 Table). The finding was supported by adding SYBR™ Green 1 nucleic acid gel stain (Invitrogen, Carlsbad, CA), where and visualized under UV light. After amplifications, positive reaction at 70 °C reactions appeared as the brightest fluorescent green under UV light by adding SYBR Green I fluorescence dye (Fig 2B). Along with the gel electrophoresis and SYBR™ Green I nucleic acid gel stain, less time was taken for amplification at that particular temperature during amplification using Genie® III. In addition to these detection methods, amplified LAMP reaction also confirmed by viewing the real-time graphs on Genie® III (Fig 2C, S2 Table).

**FIGURE 2.**
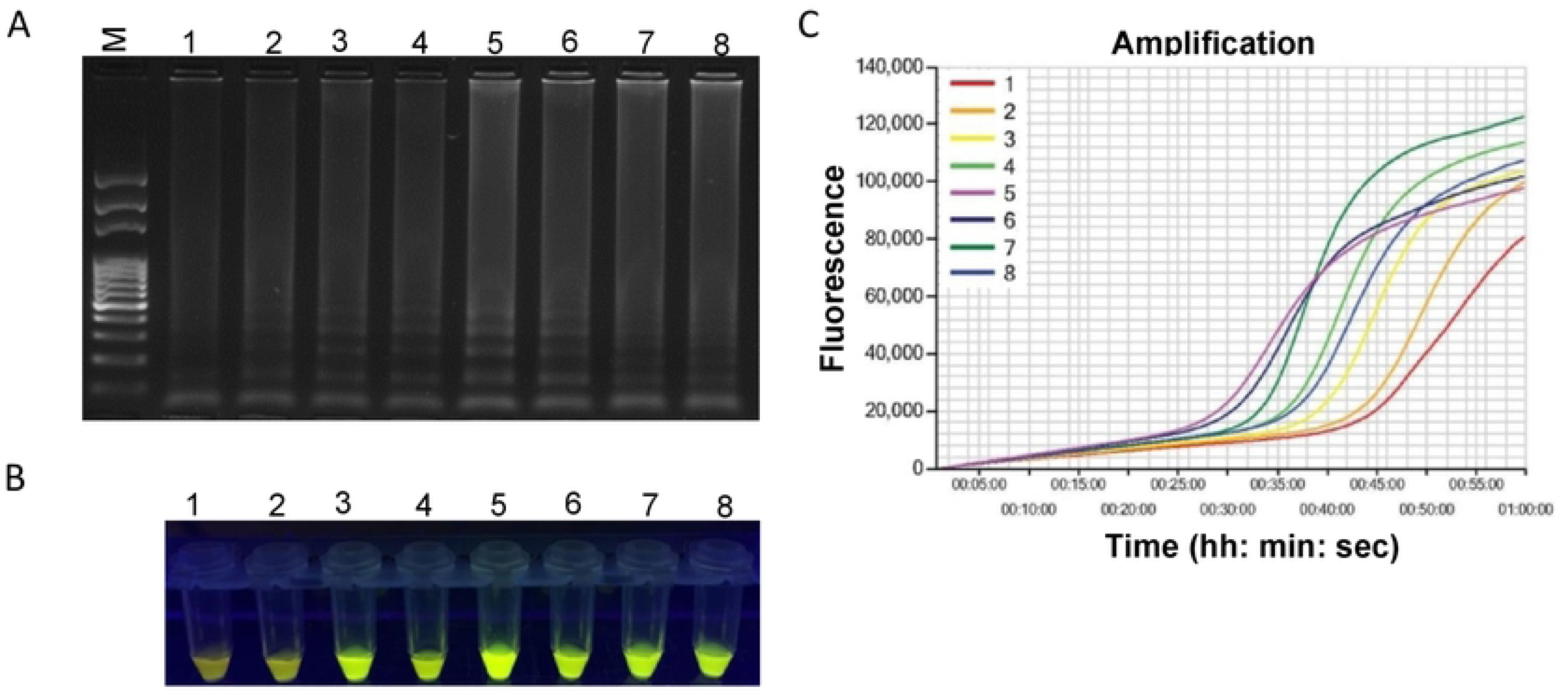
Optimization of incubation temperature for LAMP reaction. Assessment was based on (A) agarose gel electrophoresis of the LAMP products (B) visualization after addition of SYBR Green I nucleic acid stain into the reaction tubes under UV light exposure, fluorescent green color represents positive amplification; (C) real-time amplification by Genie® III. Here, 1: 66°C, 2: 67° C, 3: 68 °C, 4: 69:° C. 5: 70 °C, 6: 71°C, 7: 72°C and 8: 73°C. Lane M: 100 bp DNA ladder.

### Detection of *M. partityla* in laboratory condition

To demonstrate the applicability of the LAMP assay for the detection of *M. partityla*, the method was evaluated using DNA collected from an individual female originated from infested pecan root gall. LAMP reactions showed a positive result with the samples which were also previously identified as *M. partityla* by PCR reactions (Fig 3D) and morphological studies. All four isolates of *M. partityla* manifested positive amplification with gel electrophoresis (Fig 3A), SYBR™ Green I nucleic acid gel stain (Fig 3B) and Genie® III (Fig 3C).

**FIGURE 3.**
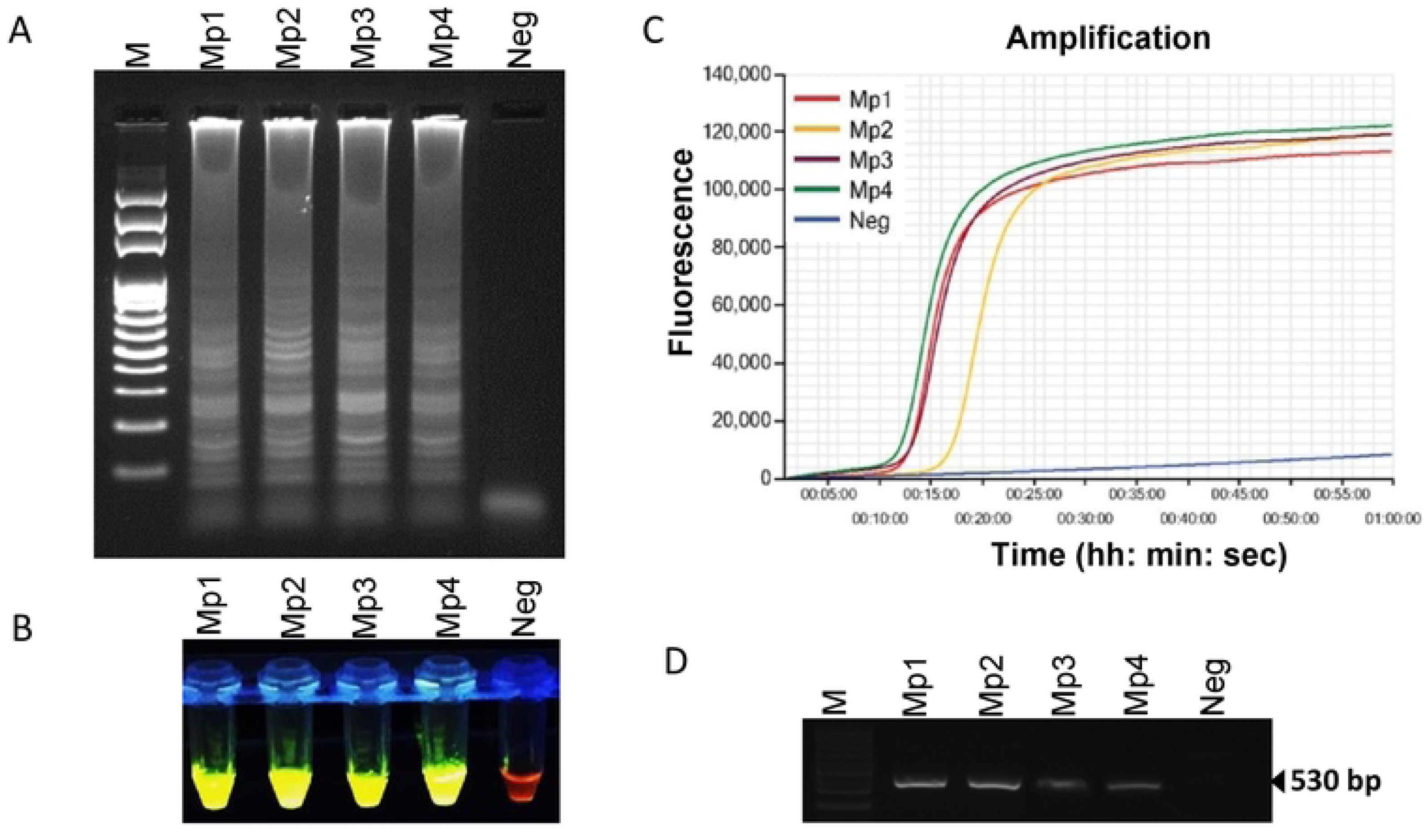
Loop-mediated isothermal amplification of DNA from four pure culture *Meloidogyne partityla.* Assessment was based on (A) agarose gel electrophoresis of the LAMP products; (B) visualization after addition of SYBR Green I nucleic acid stain into the reaction tubes under UV light exposure, fluorescent green color represents positive amplification; (C) real-time amplification by Genie® III; (D) PCR amplification using primer pair C2F3/1108. Lane M: 100 bp DNA ladder; Mp1, Mp2, Mp3 and Mp4: four different isolates of *Meloidogyne partityla*; Neg: negative control.

### Specificity of LAMP assay

Specificity of the designed primers was assessed using five *Meloidogyne* spp. Only *M. partityla* provided with positive amplification, while no amplification was observed for *M. hapla*, *M. javanica*, *M. incognita* and *M. arenaria* by the LAMP assay (Fig 4). Results were confirmed using three different detection strategies including agarose gel doc image analysis (Fig 4A), SYBR™ green based UV image (Fig 4B), Genie III amplification curve analysis (Fig 4C). All three detection strategies showed a positive reaction only from *M. partityla*, but not from other nematode species (Fig 4). The results indicated that the LAMP assay could distinguish *M. partityla*, from closely related *Meloidogyne* spp.

**FIGURE 4.**
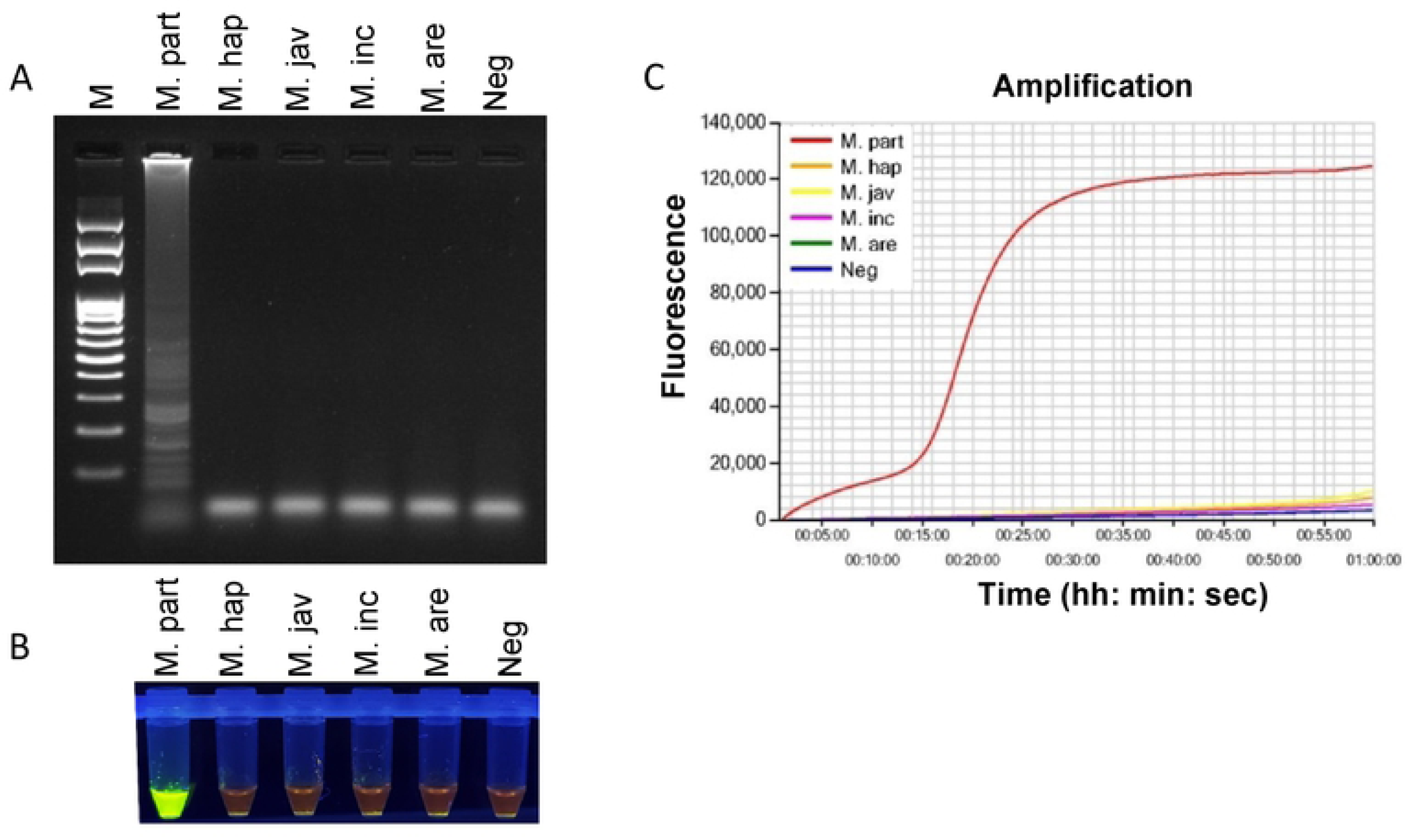
Specificity determination of LAMP assay using DNA from pure cultures of five different *Meloidogyne* spp. Assessment was based on (A) agarose gel electrophoresis of the LAMP products; (B) visualization after addition of SYBR Green I nucleic acid stain into the reaction tubes under UV light exposure, fluorescent green color represents positive amplification (C) real-time amplification by Genie III. Lane M: 100 bp DNA ladder;M. part; *Meloidogyne partityla*; M. hap: *Meloidogyne hapla*; M. jav: *Meloidogyne javanica;* M. inc: *Meloidogyne incognita*; M. are: *Meloidogyne arenaria;* Neg: negative control.

### Sensitivity comparison of LAMP with conventional PCR

Ten-fold serial dilutions of purified genomic DNA from a single female nematode (n=3) were used to evaluate the sensitivity of the LAMP method. Positive amplification were viewed with gel electrophoresis (Fig 5A), fluorescent green colors (Fig 5B), and Genie® III real-time graphs (5C) until 0.1pg per reaction, indicating that the detection limit was 1 pg of genomic DNA. For conventional PCR, expected fragment was amplified when the template DNA was diluted up to 10 pg. Thus, for purified DNA, LAMP was at least 100-fold more sensitive than conventional PCR (Fig 5D). No amplification was observed in the no-template control.

**FIGURE 5.**
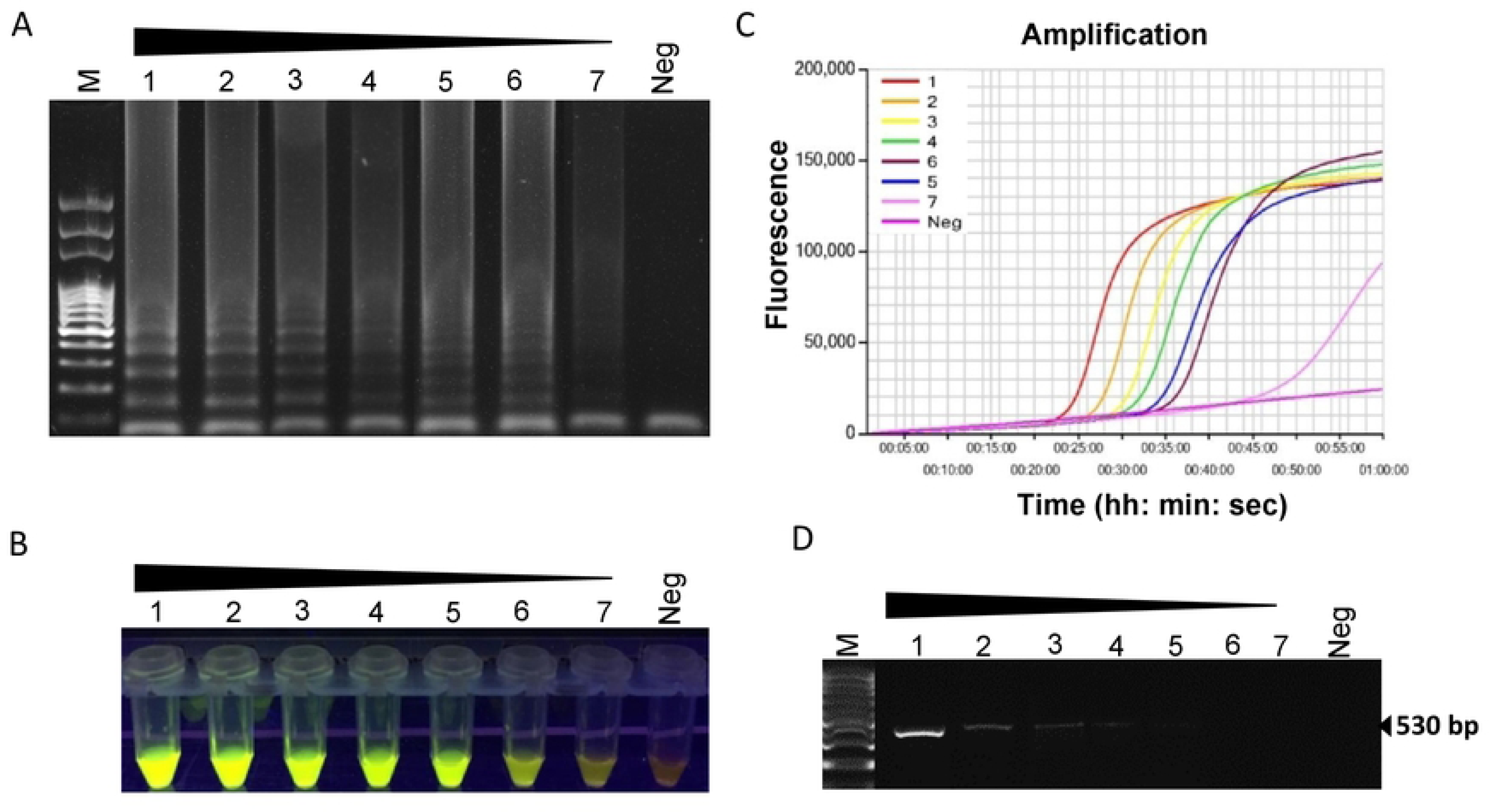
Sensitivity of LAMP assay using 10-fold serially diluted DNA extracted from pureculture of *Meloidogyne partityla*. Assessment was based on (A) agarose gel electrophoresis of the LAMP products (B) visualization after addition of SYBR Green I nucleic acid stain into the reaction tubes under UV light exposure, fluorescent green color represents positive amplification; (C) real-timeamplificationby Genie Ill (D) PCR amplification using primer pair C2F3/1108. Lane M: 100 bp DNA ladder; numbers 1 to 7: 10-fold serial dilution of *M. partityla DNA* from 100 ng/µl to 10-4 ng/µl; Neg: negative control.

### Onsite detection in infested plant root galls using LAMP assay

Seven infested root gall samples were collected from UGA Ponder research farm to evaluate the applicability of this assay for real world examples (S3 Fig). All samples showed positive to LAMP reaction confirming as *M. partityla* (Fig 6A, 6B). The results suggest that the portable handheld magnifier-based LAMP method is very useful for on-site nematode diagnosis. This also demonstrates the applicability of LAMP as a sensitive assay for determining species level at field condition.

**FIGURE 6.**
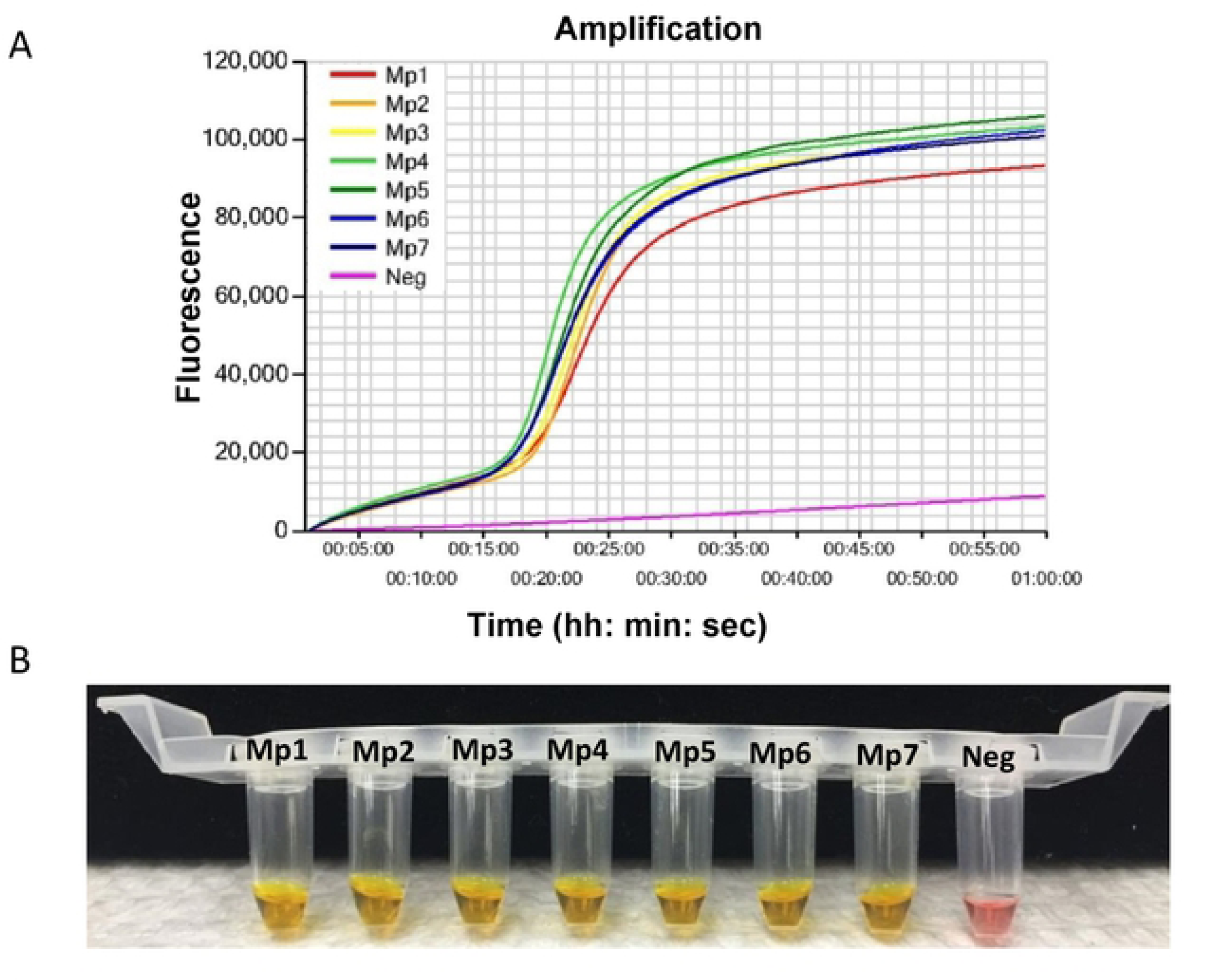
LAMP detection of *Meloidogyne partityla* from seven infected pecan root gall samples. Detection was based on (A) real-time amplification by Genie® III; (B) visualization using WarmStart colorimetric dye to determine positive reaction by naked eye for the field detection. Yellow color represents positive amplification while no amplified product remained pink. Mp1 to Mp7: seven different field isolates of *Meloidogyne* partityla Neg: negative control.

## Discussion

The specific and quick diagnosis of RKNs is required for disease prediction and adequate management control strategy selection. Traditionally *M. partityla* was diagnosed based on morphological observation, and PCR‐based molecular methods which is a time‐consuming process and requires highly traned personnel and laboratory instruments [26, 27]. The LAMP assay is an effective tool for plant-parasitic nematode identification because of its capability of DNA amplification at isothermal conditions with high sensitivity and efficiency [18, 22, 24]. In this report, we present a novel method to detect the pecan RKN, *M. partityla*. The identification can be completed in an onsite setting using a handheld magnifier and portable incubator. The amplified products can be detected visually by the naked eye and/or with the Genie® III instrument to view the amplification graph on the screen within 2 hours. Several LAMP assays have been developed to detect common *Meloidogyne* spp. [22, 24, 25]. Some of these LAMP sets are very specific to a certain RKN species, and in some cases, a single LAMP primer set can also be used for detecting double or multiple RKNs simultaneously [24, 25]. In 2019, Zhang et al. developed a LAMP primer set which can detect two potato nematodes, *M. chitwoodi* and *M. fallax*. A universal RKN-LAMP was also reported that can identify the four common *Meloidogyne* spp.: *M. incognita*, *M. arenaria*, *M. javanica* and *M. hapla* [22]. Here we designed LAMP primers targeting the conserved ITS region of *M. partityla* using the PrimerExplorer v.4 program (Fig 1). The targeted ITS region contains significant nucleotides differences between *M. partityla* and other closely related *Meloidogyne* spp. which reduces the risk of misdetection (Fig S2). Based on test results, this newly designed LAMP assay showed high specificity which only yields positive results with *M. partityla* and provides us with an essential tool to identify *M. partityla* correctly (Fig 3, 4). The results in this study exhibited high specificity to *M. partityla* populations, with no cross-reaction with other closely related RKN infested samples. Thus, the newly developed LAMP primer set could be a reliable identification tool of *M. partityla*.

Additionally, we present evidence that LAMP is faster than traditional morphology-based microscopic methods. The entire morphological diagnosis process took about 2 days which is very consistent with other reported observational methods of RKN diagnosis that mostly rely on the distinct morphological and anatomical characteristics of second-stage juveniles, adult males, as well as perineal patterns of adult females [10, 28–30]. Our results based on the morphological characteristics of J2s in this study were in agreement with those reported for *M. partityla* [26, 27] and confirms the reliability of the LAMP technique for rapid identification of *M. partityla*. PCR based traditional molecular techniques have also been frequently used to detect *M. partityla* by designing primers on the ribosomal intergenic spacer (rDNA). However, Our results showed these techniques are less sensitive than LAMP as seen by LAMP detection of 1 pg pure genomic DNA which was 100 times lower than that of PCR-based detection methods (Fig 5). This finding was supported by previously published reports on LAMP for other pathogens [21, 24, 31].

LAMP has recently been used as an on-site diagnosis technique for rapid and sensitive detection of multiple pathogens as the assay only requires a single temperature rather than a thermal cycler [32, 33]. In this study, we developed a LAMP-based quick detection protocol of *M. partityla* which can be done in on-site settings. This method can be run in the field using a colorimetric dye for visual conformation or using a real-time amplification machine for results in less than one hour [34]. In this study, direct extraction of DNA from a single female isolated from root galls greatly improved the efficiency of the Extract-N-Amp™ Tissue PCR Kit. This extraction process is rapid and only requires a portable incubator. Our onsite study demonstrated that the LAMP assay combined a handheld magnifier and an incubator was fully applicable to plant nematode detection (Fig 6, S3 Fig). To the best of our knowledge, this is the first evidence of detecting plant nematodes using the LAMP assay in a field setting. Combined with similar observations from previous reports (Niu et al., 2012, Kikuchi, 2009), these results showed that this assay has great simplicity and can overcome various types of limitations for using prepared tissue suspension, including the need for a laboratory setting. Field samples can be collected and DNA immediately extracted using the Extract-N-Amp™ Tissue PCR Kit. Those samples can then be tested for the presence of *M. partityla* using the simple, reliable LAMP method, all in the span of 60 minutes.

In summary, the portable LAMP assay described here was shown to be very reliable for the rapid detection of *M. partityla.* It is simple to operate, accurately differentiates among *Meloidogyne* species, provides results quickly, and does not require specialized equipment in comparison with the traditional morphology-based detection techniques and PCR based molecular method. The LAMP assay will be invaluable to those working with this nematode, and will greatly increase our ability to monitor and manage *M. partityla* in pecan.

## Author contributions

EA designed the project and supervised the work. SW, JB and GJ performed the experiments and analyzed the data. All authors participated in writing and editing the manuscript.

## Acknowledgments

The authors thank Latoya Hayes for technical assistance filed sampling and Owen Hudson for laboratory technical assistance.

